# The variable wheat stripe rust effector *AvrYr7* evades *Yr7* recognition through sequence and expression polymorphisms

**DOI:** 10.64898/2026.05.09.724051

**Authors:** Danish Ilyas Baig, Mareike Möller, Rita Tam, Eric C. Pereira, Julian Rodriguez-Algaba, Shideh Mojerlou, Mogens Støvring Hovmøller, Annemarie Fejer Justesen, Thuan Nha Ho, Jinwen Zhang, Yi Ding, Jingyu Li, Jiajie Wu, Sambasivam Periyannan, Xiaoxiao Zhang, John P. Rathjen, Benjamin Schwessinger

## Abstract

Wheat provides about 20% of total dietary calories worldside^1^. Wheat diseases, including wheat stripe (yellow) rust, cause billions of dollars in losses each year^2^. Wheat stripe (yellow) rust is caused by the fungal pathogen *Puccinia striiformis* f. sp. *tritici* (Pst) which is best controlled by fungicide application and disease resistant wheat cultivars^3^. To-date, there are over 80 catalogued and >10 cloned yellow rust resistance genes (*Yr* genes)^4^. Yet our knowledge of corresponding avirulence (*Avr*) genes lags far behind^5–8^. The absence of cloned *Avrs* reflects *Pst’s* complex genome and the lack of robust transformation and genetic systems^3^. Recent advances in generating high-quality genome assemblies and the development of wheat defense assays have addressed these challenges^9–11^. Here we clone AvrYr7 which is recognized by Yr7^12^. We further identify six additional alleles of *AvrYr7* that escape recognition due to non-synonymous genetic variations, transposable element activity, missense mutation, and expression polymorphism. These findings provide critical insights into virulence evolution in one of the world’s most important wheat pathogens.

## Main text

Wheat yellow rust resistance (*Yr*) genes are often classified into either all-stage resistance that is effective throughout plant development or adult-plant resistance that is typically quantitative and expressed at later stages of wheat development^4^. Most all-stage *Yr* resistance genes fall into the NBS-LRR (nucleotide binding site – leucine rich repeats) *R* gene class and are predicted to recognize specific *Pst* effectors in the wheat cytoplasm^4,13^. Effector alleles, which protein products are recognized by *R* genes, are called Avirulence genes (*Avr*) because they render *Pst* isolates avirulent on wheat cultivars that carry the corresponding *Yr* gene. All-stage *Yr* genes often become ineffective after deployment in agriculture due to the adaptation of *Pst* populations via variations in corresponding *Avr* alleles causing loss of recognition^3,14^. Genetic diversity in *Pst* arises through multiple evolutionary routes, including accumulation of mutations during long-term asexual reproduction; rare, geographically restricted sexual recombination; and somatic hybridisation^3,15,16^. The latter is facilitated by *Pst’s* genome organization as a dikaryon that partitions its two genome copies into separate nuclei within the same cytoplasm^3^. No *Pst Avr* locus has been identified to date, and we lack direct examples of virulence evolution caused by genetic variation at *Pst* avirulence loci.

*Yr7* was one of the first all-stage *Yr* resistance genes to be cloned. It encodes a NBS-LRR immune receptor with a non-canonical zinc-finger BED domain N-terminal to the classical CC (coiled coil) domain^12,17,18^. To identify the corresponding *Pst* avirulence effector, we build on our recent high-quality reference genome of an isolate (Pst104E) from the clonal PstS0 lineage^9^.

Avirulence to *Yr7* appears to be ancestral in PstS0, as most isolates including Pst104E, DK214_12, and the earliest sampled isolates are avirulent^19,20^. We analyzed two virulent isolates from this lineage, NL72067 and DK06_00, that cause disease on the *Yr7* differential wheat cultivar Lee (Fig. 1a)^21^. To identify candidate effector genes that might confer avirulence to *Yr7*, we sequenced the genomes of the two *Yr7* virulent isolates NL72067 and DK06_00, that were collected 28 years apart in the Netherlands (1972) and in Denmark (2000), respectively. We hypothesized that the *Avr* gene would most likely be heterozygous in Pst104E (1982), while only the recognized allele would be mutated in the two *Yr7* virulent isolates NL72067 and DK06_00. We base our hypothesis on the assumption that in a dikaryon during clonal reproduction, it is very unlikely that two independent mutations render both alleles of a homozygous *Avr* locus non-functional at the same time in absence of recombination or gene conversion^22,23^. Using Pst104E as a *Yr7* avirulent reference, we identified non-synonymous and missense mutations in heterozygous candidate effector genes^9^. We identified a total of 51 candidate effector genes with mutations in both virulent isolates compared to the reference (Table S1). The gene *Pst104Ev4_29515* caught our attention, because it contained independent mutations in NL72067 and DK06_00 (Fig. 1b). In addition, *Pst104Ev4_29515* is located on the left arm of chromosome 7, which is consistent with previous studies that mapped an avirulence gene cluster, including *AvrYr7, AvrYr44, AvrYr43, AvrYrExp2*, to a similar chromosomal location^22,23^. *Pst104Ev4_29515* encodes a 430 amino acid (aa) protein with unknown function. In DK06_00, a missense mutation results in a premature stop codon at position 260, while in NL72067 a 179 bp insertion of terminal inverted repeat DNA transposon (Tc1/Mariner-like) causes a frameshift and altered C-terminal amino acid sequence starting at position 389. The alternate allele, *Pst104Ev4_08667*, is identical in all three isolates but differs from *Pst104Ev4_29515*. It is 442 aa long, has a K->E substitution at position 233, and a frameshift altering the C-terminal tail from position 414 (Fig. 1c). We hypothesized that *Pst104Ev4_29515* is the allele recognized by *Yr7* in Pst104E, based on the identification of independent mutations in the two *Yr7* virulent isolates and the presence of an alternate allele that is sequence conserved in all three isolates. Based on these findings, we designated *Pst104Ev4_29515* as *AvrYr7-01, Pst104Ev4_08667* as *avrYr7-02*, and mutant alleles of *AvrYr7-01* in DK06_00 and NL72067 as *avrYr7-03* and *avrYr7-04*, respectively.

**Figure 1:**
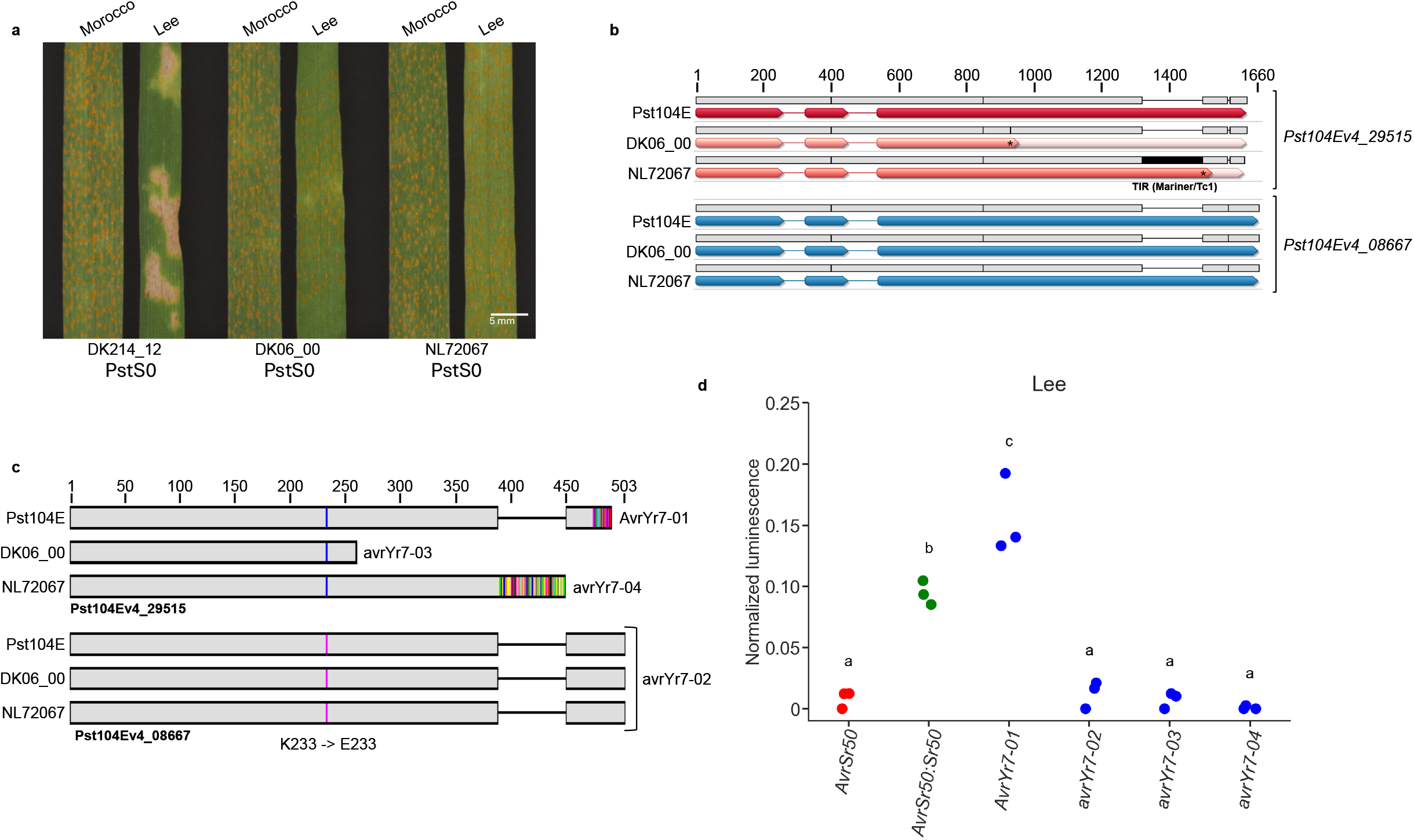
Genetic variation within the PstS0 lineages identifies *AvrYr7*. a) Representative infection phenotypes of *Puccinia striiformis* f. sp. *tritici* (*Pst*) isolates from the clonal *PstS0* lineage on cv. Morocco and cv. Lee (*Yr7*). Leaves of wheat cv. Morocco (susceptible control) and cv. Lee carrying the resistance gene *Yr7* were inoculated with the *Pst* isolates DK214_12, DK06_00, and NL72067. Infection types (ITs) were scored visually on the second leaves 18 days after inoculation using a 0–9 scale, where ITs 0–6 were classified incompatible and 7–9 compatible at the phenotypic level^26,52,53^. All three isolates produced abundant uredinia on Morocco (IT 7-8). On cv. Lee, DK214_12 induced an incompatible chlorotic/necrotic response with little or no sporulation (IT 2-3), whereas DK06_00 and NL72067 produced compatible phenotypes with abundant uredinia (IT 7). Scale bar = 5 mm. b) Alignment of genes encoding for Pst104Ev4_29515 and Pst104Ev4_08667 in the avirulent PstS0 isolate 104E and the two virulent PstS0 isolates DK06_00 and NL72067. Exons are indicated as arrows and introns as thin lines connecting the exons. Black lines indicate differences between alignments. Pst104Ev4_29515 differs between all three isolates, with a point mutation in the third exon resulting in a premature stop in DK06_00 and a 179bp insertion close to the C-terminus in a frameshift in NL72076. Pst104Ev4_08667 is identical in all three isolates. c) Protein alignment of all four alleles found in the three PstS0 isolates. AvrYr7-01 (Pst104Ev4_29515 in 104E), avrYr7-02 (Pst104Ev4_08667), avrYr7-03 (Pst104Ev4_29515 in DK06_00), avrYr7-04 (Pst104Ev4_29515 in NL72076) highlighting mutations and differences in the C-terminal region. d) Defense reporter activation of *AvrYr7-01* in cv. Lee. *AvrYr7-01* activated the defense signaling in protoplasts of a *Yr7* containing wheat cv. Lee while no defense activation by the alternate allele (*avrYr7-02*) and the PstS0 mutant alleles (*avrYr7-03* and *avrYr7-04*). *AvrSr50* and *AvrSr50:Sr50* served as negative and positive controls, respectively.

We made use of our recently established wheat protoplast defense reporter assay where *Avr-R* recognition triggers the quantitative expression of luciferase regulated by defense response promoters and where the normalized luciferase activity is used as read-out for wheat defense activation^10^. We applied this wheat protoplast defense reporter assay to cv. Lee (*Yr7)* to test our hypothesis that *AvrYr7-01* is the recognized allele and *avrYr7-02, avrYr7-03* and *avrYr7-04* are non-recognized alleles. Expression of *AvrYr7-01* in cv. Lee induced the defense reporter comparable to the positive control of the co-expressed *Avr-R* gene pair *AvrSr50-Sr50* (Fig. 1d). In contrast, *avrYr7-02, avrYr7-03* and *avrYr7-04* did not induce the defense reporter with normalized luminescence levels similar to that of the non-recognized effector *AvrSr50* (Fig.1d)^24^. Classification of *AvrYr7-01* as the avirulent recognized allele and *avrYr7-02* as the virulent non-recognized allele was further validated by defense reporter induction by *AvrYr7-01*, but not *avrYr7-02*, in protoplasts of the wheat cultivars Thatcher and Vixen, both of which encode *Yr7* (Extended Data Fig. 1)^12,25^. We further validated the *Yr7* specificity by expressing *AvrYr7-01* and *avrYr7-02* in protoplasts derived from *Triticum spelta* encoding the *Yr7*-related *Yr5*^12^ and the susceptible wheat cultivars Morocco and Avocet S, that do not contain any major *Yr* gene including *Yr7*^12,26^. Neither *AvrYr7-01* nor *avrYr7-02* induced defense reporters when expressed in these protoplasts (Extended Data Fig. 2).

We designed an experiment to test for recognition specificity of *AvrYr7-01* by *Yr7* and to rule out recognition by shared yet undescribed *Yr* genes encoded in cv. Lee, Thatcher, and Vixen. We co-expressed *AvrYr7-01* or *avrYr7-02* in the presence and absence of *Yr7* in protoplasts derived from cv. Avocet S. We did not observe defense reporter activation when expressing either allele in cv. Avocet S in the absence of *Yr7*, or when expressing *Yr7* alone. Similarly, the defense reporter was not activated when co-expressing *Yr7* with the unrelated effector *AvrSr50* or with the non-recognized allele *avrYr7-02* (Fig. 2a, Extended Data Fig. 2). Strong defense reporter activation was detected only when *Yr7* was co-expressed with *AvrYr7-01*, demonstrating the specificity of the *AvrYr7-01-Yr7* interaction.

**Figure 2:**
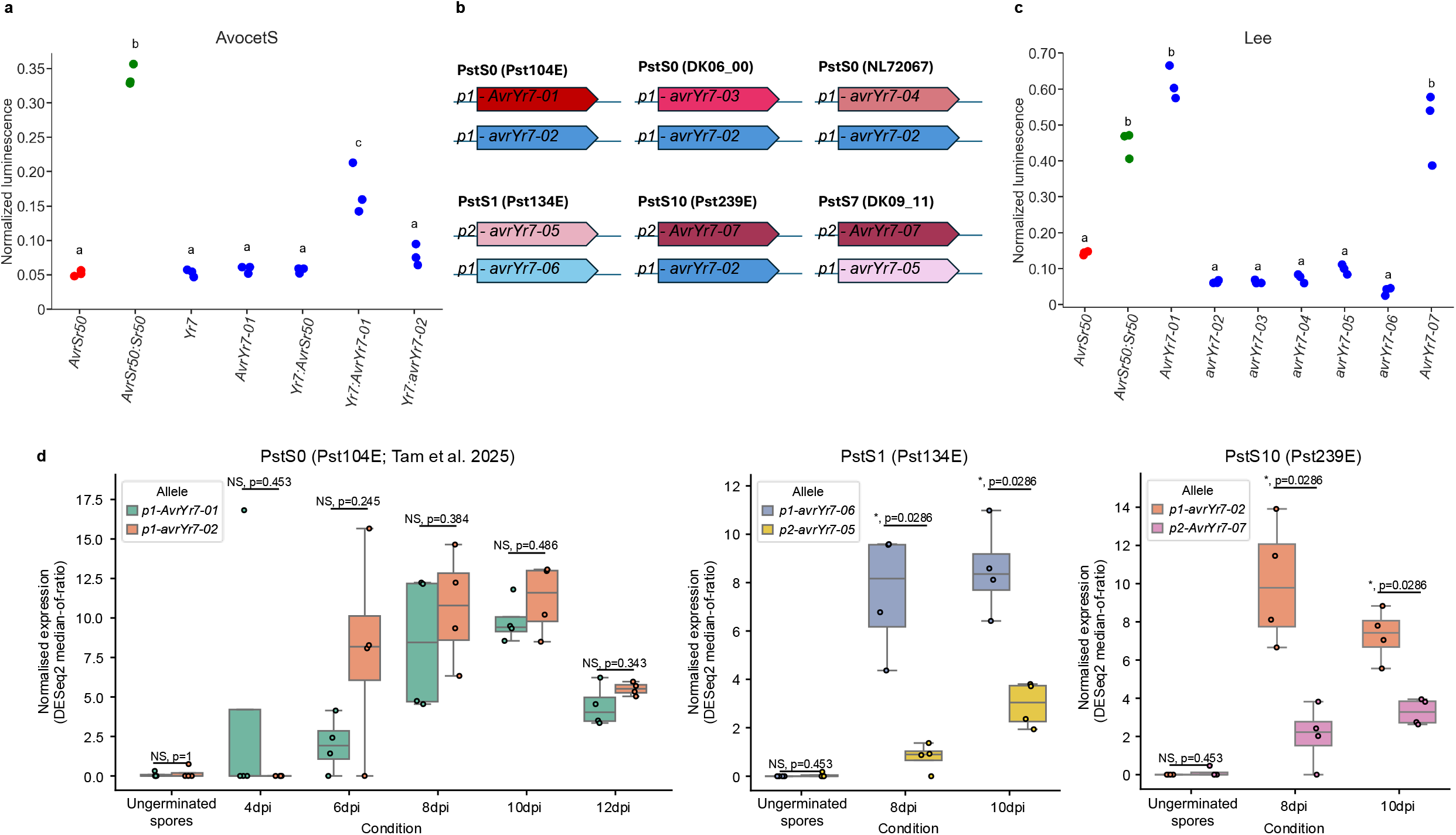
AvrYr7 protein sequence variation and expression polymorphisms lead to escape of Yr7 recognition. a) Defense reporter activation of *AvrYr7-01* with *Yr7* in AvocetS. Co-expression of *AvrYr7-01* with *Yr7* activated the defense signaling in protoplasts of a non-*Yr7* wheat cv. Avocet S while no defense activation by *avrYr7-02* with *Yr7. AvrSr50* and *AvrSr50:Sr50* served as negative and positive controls, respectively. b) Schematic representation of avrYr7 alleles detected in six different Pst isolates. We found a total of seven different alleles and unique allele combinations in each isolate. c) Defense reporter activation of *AvrYr7-01* and *AvrYr7-07* in Lee. While lining up all the various alleles of *AvrYr7* in protoplasts of a *Yr7* containing wheat cv. Lee, defense signaling was activated by *AvrYr7-01* and no defense activation by any other allele except *AvrYr7-07. AvrSr50* and *AvrSr50:Sr50* served as negative and positive controls, respectively. d) Expression profiles of *AvrYr7* alleles in spores and during infection on the susceptible cv. Morocco in isolates Pst104E (PstS0, avirulent on cv. Lee), Pst134E (PstS1, virulent on cv. Lee) and Pst239E (PstS10, virulent on cv. Lee). Shown are normalized counts for four replicates per condition. *AvrYr7* alleles are not expressed in ungerminated spores. In Pst104E, both alleles, the recognised *p1-AvrYr7-01* and the non-recognised *p1-avrYr-02* show similar expression levels throughout infection. In contrast, alleles *p2-avrYr7-05* in Pst134E and *p2-AvrYr7-07* in Pst239E are significantly lower expressed as their alternate allele.

Most current major *Pst* lineages are virulent on *Yr7* (Extended Data Fig. 3)^27^, which encouraged us to explore the genetic variation of *AvrYr7* alleles in globally distributed *Pst* lineages^28^. We identified and characterized alleles in isolates from the *PstS1* (Pst134E), *PstS10* (Pst239E) and *PstS7* (DK09_11) lineages^16^. We identified three additional alleles of *AvrYr7* (*avrYr7-05, avrYr7-06* and A*vrYr7-07*), containing non-synonymous mutations compared to the previously identified alleles *01* to *04* (Fig. 2b, Extended Data Fig. 4, Table S2). Considering all these isolates are virulent on cv. Lee (*Yr7)*, we hypothesized that none of these alleles would induce a defense response in the presence of *Yr7*. Consistently, *avrYr7-05* and *avrYr7-06* did not induce defense reporter activation in cv. Lee derived protoplasts (Fig. 2c). In contrast, *AvrYr7-07* induced defense reporter activation to a level similar to that of *AvrYr7-01*, despite *AvrYr7-07* being encoded in the genomes of PstS10 (Pst239E) and PstS7 (DK09_11), both of which are virulent on wheat cultivars carrying *Yr7*^27,28^.

We explored three independent hypotheses for why the recognition of *AvrYr7-07* in *Yr7* expressing protoplasts did not translate to avirulent phenotypes on cv. Lee for Pst239E or DK09_11. First, the alternate *avrYr7* allele might suppress the function of *AvrYr7-07 in planta* as a dominant negative allele^29^; Second, *AvrYr7-07* might evade recognition due to a lower sensitivity of *Yr7* when compared to *AvrYr7-01*^6,30^; Third, *AvrYr7-07* might be expressed at a level too low for functional recognition by *Yr7*^30–34^.

To test the first hypothesis, we co-expressed *AvrYr7-01* and *avrYr7-02* (the allele combination in avirulent isolate Pst104E) and *AvrYr7-07* and *avrYr7-02* (the allele combination in virulent isolate Pst239E) in cv. Lee protoplasts. Both allele combinations induced defense reporter activation, as did *AvrYr7-01* and *AvrYr7-07* alone, but not *avrYr7-02* (Extended Data Fig. 5). These results did not support the hypothesis that a paired alternate non-recognized allele could suppress the recognition of *AvrYr7-07* by *Yr7 in planta*.

To test our second hypothesis, we varied the plasmid amounts used to express *AvrYr7-01* and *AvrYr7-07* in cv. Lee protoplasts because we previously showed that defense reporter activation is dosage dependent^10^. If Yr7 were to display a reduced sensitivity to AvrYr7-07 we would expect to see reduced or no defense activation at lower plasmid amounts when compared to AvrYr7-01. This was not the case, as both *AvrYr7-01* and *AvrYr7-07* induced comparable defense reporter activation at all plasmid amounts with a slight but not statistically significant reduction on the level of defense reporter activation at lower plasmid doses (Extended Data Fig. 6).

To investigate if hypoexpression of *AvrYr7-07* might explain the *Yr7* virulence phenotype of Pst239E and DK09_11, we compared the expression level of *AvrYr7-07* with five other alleles in three different isolates. The presence of diagnostic SNPs, as well as long-read RNA-seq data, enabled us to differentiate the expression levels of the two alleles in each *Pst* isolate^9^.None of the *AvrYr7* alleles were expressed in spores and expression was only detected *in planta*, supporting their roles during the biotrophic wheat infection phase (Fig. 2d). For Pst104E, *AvrYr7-01* and *avrYr7-02* were expressed at comparable levels throughout the infection time course. In Pst239E, *AvrYr7-07* was significantly lower expressed than *avrYr7-02*. In Pst134E, *avrYr7-05* was significantly lower expressed than *avrYr7-06* (Fig. 2d, Extended Data TableS3). We investigated the 1 kb upstream putative promoter regions of *AvrYr7-07* and *avrYr7-05* to identify genetic variations that might explain the observed expression polymorphism. Indeed, both *AvrYr7-07* and *avrYr7-05* share two base pair substitutions at -623 (G to T) and -882 (C to T) whereas all other positions were fully conserved between all alleles (Extended Data Fig. 7). We designed the putative promoter sequence identical to that of *AvrYr7-01* as *p1* and *p2* in the case of *AvrYr7-07* (Fig. 2b and Extended Data Fig. 7). We therefore hypothesize that these shared upstream substitutions in *p2* lead to reduced expression of *AvrYr7-07* and *avrYr7-05*, and that in the case of *AvrYr7-07* the reduced expression underlies the virulent phenotypes of Pst239E and DK09_11 on *Yr7* wheat cultivars. We hence extended the recently introduced *Avr* allele naming convention for cereal rusts^6^ to include changes in regulator promotor sequences with a demonstrated impact on avirulence phenotypes (see Methods Section for details) (Fig. 2b).

Lastly, we explored the sequence and predicted structural differences between the AvrYr7 protein variants, including potential interaction interfaces with Yr7^12,35^. The aa multiple sequence alignment showed high sequence conservation among AvrYr7 variants except for the C-terminal region equivalent to AvrYr7-01 residues 414-430 (Fig. 3a). Compared to AvrYr7-01, avrYr7-03 is truncated at residue 259 and avrYr7-04 is variable from residue 389. Notably, AvrYr7-01 and avrYr7-02 differ by only one amino acid at residue 233 (K→E) in the sequence conserved N-terminal region (AvrYr7-01 1-413 aa). This K→E substitution is shared with another non-recognized variant avrYr7-06, which differs from avrYr7-02 by only a single amino acid in the predicted signal peptide at residue 8 (L→I). The unrecognized avrYr7-05 variant has two amino acid differences compared to AvrYr7-01; one in the conserved region at 229 (I→T) and one in the C-terminal region at 423 (A→T). The latter substitutions is also present in the (in protoplast-recognized) variant AvrYr7-07 suggesting that the 229 (I→T) substitution mediates the loss of recognition of avrYr7-05.

**Figure 3:**
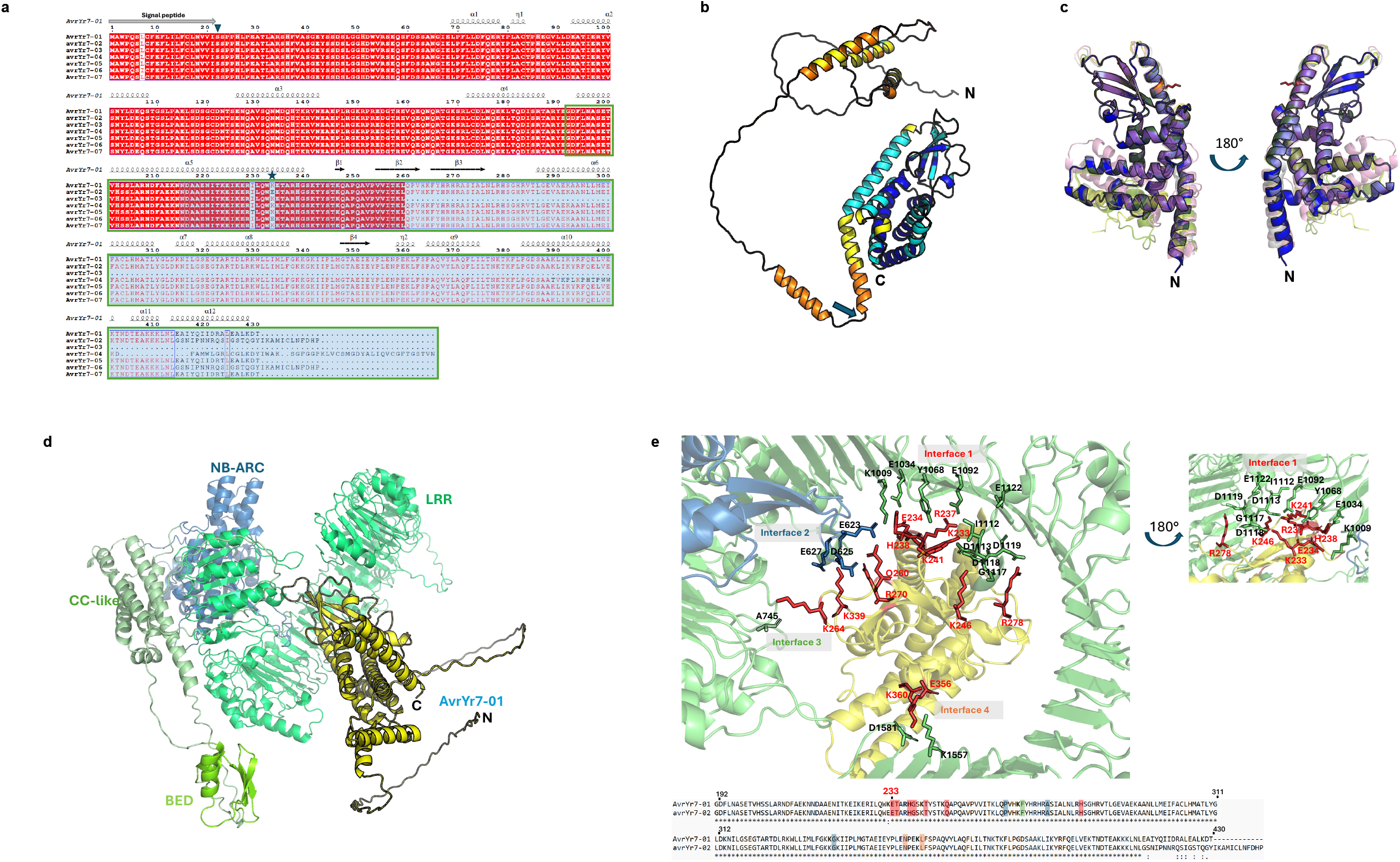
Structural models of AvrYr7 variants and its interaction complex with Yr7 provide hypotheses for loss of recognition. a) Multiple sequence alignment of AvrYr7 variants. The residues are numbered according to AvrYr7-01. The secondary structure elements of the Alphafold3 predicted AvrYr7-01 model are displayed on the alignment. The N-termini of the mature protein sequences used for AlphaFold3 modelling is indicated by a triangle. The region with high structural similarities starts at residue 192, and the region with high modelling confidence scores starts at residue 215, indicated by green frames and blue boxes, respectively. The key variable residue 233 is indicated by a star. b) Structure model of AvrYr7-01 mature protein without signal peptide, coloured according to Alphafold3 modelling confidence scores: dark blue, very high; light blue, confident; yellow, low; orange, very low. The arrow marks the beginning of the C-terminal region used for structural alignment in (c). c) Superimposition equivalent sequences to AvrYr7-01 192-430 aa of six AvrYr7 variants, excluding avrYr7-03. AvrYr7-01 is coloured in blue and outlined. Other variants are coloured as green, avrYr7-02; pink, avrYr7-04; grey, avrYr7-05; yellow, avrYr7-06; salmon, AvrYr7-07. Key variable residue K233 of AvrYr7-01 is shown as sticks and coloured in red. d) Structural model of AvrYr7 and Yr7 interaction. AvrYr7-01 is coloured in yellow. Yr7 is coloured and labelled according to its sub-domains. e) Predicted interaction interface between AvrYr7 and Yr7. Upper panel, the interface residues are shown as sticks and coloured in red for AvrYr7-01 and blue or green for Yr7 as in (**d**). The Yr7 and AvrYr7-01 interface residues are labelled in black and red respectively. Lower panel, pairwise sequence alignment of AvrYr7-01 and avrYr7-02 with key residues highlighted in red, blue, green and orange for interface 1-4 respectively.

AlphaFold3^35^ modelling of the mature AvrYr7-01 predicted structural features with high confidence starting from 215 aa till the C-terminus at 430 aa (Fig. 3b). Conserved domain and structural homology searches did not identify any homologs. Structural alignment among AvrYr7 variants revealed a variable N-terminal region between residues 22-191, which may be caused by the low model confidence. Despite the amino acid sequence variation in the C-terminal tails between AvrYr7 variants, the region between residues 192-430 forms a core structure highly similar to AvrYr7-01, except for avrYr7-03 which lacks the C-terminal 171 aa (Fig. 3c). The two key amino acids I229 and K233, which substitutions likely define non-recognition in avrYr7-02, -05, and -06, are both located on α5-helix (Fig. 3a-c). While K233 is predicted to be surface exposed, I229 is buried between the α5- and α6-helix and its substitution to T leads to a subtle predicted conformational change of the α5-helix. Indeed, the C-terminal end of α5-helix is predicted to be a main interaction interface between AvrYr7-01 and Yr7 where K233 forms direct interaction with the Yr7 LRR domain (Fig. 3d&e). These structural models provide a rational hypothesis for the role of I229 and K233 in mediating Yr7 recognition and their substitutions in AvrYr7 variants loss thereof. Future experiments will have to explore if AvrY7-01 directly binds to Yr7 and if these predicted interaction surfaces truly inform the AvrY7-Yr7 recognition spectrum.

Our analysis identifies *AvrYr7* as the first *R* gene matched avivrulence effector for *Pst*, where the avirulence causing allele *AvrYr7-01* is recognized by *Yr7* specifically. This contrasts with *AvrPstB48* which elicits defense reporter activation in a defined subset of wheat cultivars but could not be matched to a cloned *Yr* gene^36^. Hence, our work closes the gap between *Pst* and all other major biotrophic and hemibiotrophic wheat pathogen like *P. graminis* f. sp. *tritici, P. triticina, Blumeria graminis* f. sp. *tritici*, and *Zymospetoria tritici* for which several *Avrs* have been identified^5–8^. In addition to the identification of *AvrYr7*, we also provide the first molecular insight into virulence evolution in *Pst*. It appears that virulence to *Yr7* in *Pst* is achieved via three distinct molecular and evolutionary mechanisms including step-mutations in the asexual lineage PstS0, expression polymorphisms, and standing genetic variation within the global pathogen population.

Step-mutations that lead to inactivation of avirluence genes are common in plant pathogens due to the extensive selection pressure imposed by the deployment of the same *R* gene across large growing areas^3,37,38^. These have been long hypothesized to contribute to virulence evolution in *Pst*, especially during its clonal reproduction stage^3,39^. We observe two independent mutational events in the *Yr7* virulent isolates belonging to the long-term clonal PstS0 lineage. A single point mutation leads to an early stop codon and a truncated version of AvrYr7 in DK06_00. In NL72067 we identified an insertion of a Tc1/Mariner-like transposable element that leads to a frameshift changing the C-terminal end of the encoding protein. The Tc1/Mariner-like insertion displays features of a completely conserved terminal repeat indicating very recent transposon activity^40^. Transposons are significant drivers of pathogen evolution including their demonstrated role of avriluence gene disruption in cereal rust fungi^41,42^. Further studies will have to explore the scale of transposon activity in *Pst* with a focus on its evolution and host adaptation during clonal reproduction.

Expression polymorphisms and gene silencing can also lead to the evolution of virulence in plant pathogens, but is a less well described phenomenon^37^. Here we show that two substitutions in the putative promoter region of *AvrYr7*, referred to as *p2*, lead to a significantly reduced mRNA expression level of alleles regulated by *p2* vs. *p1*. The reduced expression of *AvrYr07-7* appears to be causative for the *Yr7* virulence phenotype of *Pst* isolates Pst239E and DK09_11, because the AvrYr7-07 protein variant is still recognized in wheat protoplast. This is consistent with observations made for the oomycete pathogen *Phytophthora sojae* where gene silencing of *Avr3a* and *Avr1b* led to virulence on hosts carrying the corresponding *R* gene^33,34^. In the case of *Pst*, the unmatched avirulence effector candidate *PST130_P495001* was proposed to contribute to virulence evolution in the PstS10 lineage via hypoexpression^31^.

Similar observations have been made for *AvrRp1-D* in *P. sorghi*^32^ and *AvrSr27* alleles in *P. graminis* f. sp. *tritici*^30^. Future experiments will have to explore the extent to which expression polymorphisms contribute to virulence evolution in *Pst* and other cereal rust fungi more broadly.

Standing genetic variation at the *AvrY7* loci also appears to contribute to *Yr7* virulence evolution in agricultural settings. Protein variants encoded by the alleles *avrYr7-02* and *avrYr7-05* are clearly different from AvrYr7-01, avrYr7-05, and AvrYr7-07 with the former having distinct C-terminal sequences starting from residue 414 aa, in addition to the more central K233E substitution. This indicates potential distinct evolutionary histories of the various *AvrYr7* alleles which might be linked to distinct nuclear haplotype evolution in *Pst*^43,44^. Future experiments will have to explore effector evolution in the proposed distinct nuclear haplotype populations of *Pst*^42,43^ and how somatic hybridization contributes to virulence evolution in agricultural settings^16^.

## Methods

### Fungal isolate description and Infection conditions

Selected *Pst* lineage:isolates (PstS0: Pst104E, DK214_12, DK06_00, and NL72067; PstS1: Pst134E and RW10_15; PstS7: DK09_11; PstS8: DK02d_12; PstS10: DK267_17 and Pst239;PstS13:DK69_15) were inoculated on wheat cv. Lee carrying *Yr7* and on the susceptible control cv. Morocco^20,27,39,45–50^. For each host genotype, 8 seeds were sown in 10-cm pots containing peat-based substrate and maintained under controlled greenhouse conditions (18□°C/12□°C day/night, 70–80% relative humidity, 16-h photoperiod). Seedlings were inoculated at the one-leaf stage, when the first leaf was fully expanded and the second leaf was half-emerged (10–12 d after sowing). For each isolate, about 20 mg urediniospores were suspended in Novec™ 7100 (3M) and applied evenly using an airbrush spray gun^26,51^. After inoculation, plants were incubated for 24 h at 12°C in darkness under 100% relative humidity and then transferred to spore-proof greenhouse cabins under the same conditions described above. Infection types (ITs) were scored visually 18 d after inoculation using a 0–9 scale^26,52,53^, where ITs 0–6 were classified as incompatible (resistant) and 7–9 as compatible (susceptible). Representative images were captured at the same time point.

### DNA extraction and long-read sequencing

High-molecular-weight genomic DNA of PstS0 isolates DK06 and NL720 and was extracted from urediniospores as described here^54,55^. DNA was size-selected and further cleaned up with a 5% PVP size selection buffer^56^. Nanopore sequencing library was prepared using the Ligation Sequencing Kit v14 (SQK-LSK114 V14), then sequenced on a P2 Solo sequencer using FLO-PRO114M flowcells to achieve ∼60x coverage.

### Genome assemblies

Reads were basecalled with dorado (v0.7.2) and the dna_r10.4.1_e8.2_400bps_sup@v5.0.0 model. Reads were subsequently error corrected using herro (v1)^57^ and both raw and error corrected reads used as input for genome assembly using verkko v2.0^58^. Genome assemblies were scaffolded with RagTag^59^ using the Pst104E genome as reference^9^. After scaffolding both assemblies contained 36 chromosome scale contigs including telomeres on both ends of each chromosome.

### Identification of *AvrYr7* candidate gene within *PstS0* isolates

The genome assemblies of the two PstS0 isolates virulent on Yr7 (DK06_00 and NL72067) were aligned independently against the reference genome of the *Yr7* avirulent PstS0 isolate Pst104E^9^ using minimap2^60^ in assembly mode. Pairwise alignments in PAF format were used for variant calling with paftools. Variant calls were intersected with the CDS coordinates of predicted 104E secretome genes using bedtools^61^ to identify genes carrying sequence variation relative to the reference. To assess whether variants were predicted to alter coding sequence, variants were annotated with bcftools csq^62^ using the 104E reference genome. Shared mutated secretome genes between DK06_00 and NL72067were identified, and only heterozygous genes with non-synonymous variants in both assemblies were retained and considered for further analyses. The 179 bp insertion in NL72067 was blasted against the RepBase^63^ 31.04 repeat library for fungi to check whether it was derived from transposable element.

### Cloning for protoplast expression vectors

Synthetic double-stranded DNA fragments (gBlocks; Integrated DNA Technologies, USA) encoding the *Pst AvrYr7-01* (*Pst*104Ev4_29515), its alternate allele (*avrYr7-02*), mutant forms (*avrYr7-03* and *avrYr7-04*) and allelic variants in different *Pst* lineages (*avrYr7-05, avrYr7-06*, and *AvrYr7-07*) were synthesized with flanking BsaI recognition sites and 5’ AATG and 3’ GCTT overhangs compatible with Level 1 Golden Gate assembly^64^. Following BsaI digestion, inserts were ligated into BsaI-linearized pWDV1 vector^10^ using 25 cycles of 37 °C (1.5 min) and 16 °C (3 min). Assembly reactions were incubated at 50 °C (5 min) and 80 °C (10 min) to inactivate the enzyme, then transformed into chemically competent NEB 5-alpha *Escherichia coli* cells. Cells were incubated on ice for 30 min, heat shocked at 42 °C for 30 s, and chilled on ice for 5 min. Then cells were recovered in SOC medium (200 μL; containing 2% tryptone, 0.5% yeast extract, 10 mM NaCl, 2.5 mM KCl, 10 mM MgCl□, 10 mM MgSO□, and 20 mM glucose) at 37 °C for 1 h with shaking (280 rpm) and plated on LB agar containing 35 µg/mL chloramphenicol. Colonies were screened by colony PCR, and positive clones were cultured for plasmid extraction (Qiagen Miniprep). Construct integrity was initially assessed by diagnostic restriction digestion and subsequently confirmed by whole-plasmid Oxford Nanopore (ONT MinION) sequencing. Reads were aligned to the reference plasmid in Geneious Prime v2025.1.3 using Minimap2^60^ to verify insert orientation and sequence accuracy. All constructs used and generated in this study are listed in TableS4.

### Defense reporter assays in wheat protoplasts

Plasmid DNA (1 μg μL^−1^) was prepared using SV Wizard mini-prep and maxi-prep kits (Promega). Protoplast isolation, PEG-mediated transfection and dual luciferase reporter assay following Wilson et al. (2024)^10^ was adapted to a 96-well format^65,66^ with further optimization^67^. All *Avr, avr, R*, and luciferase reporter constructs were based on the self-replicating pWDV1 vector and driven by the maize ubiquitin (UBI) promoter^10^. Per replicate, 1 μg *Avr/avr*, 2 μg *R*, and 5 μg luciferase reporter plasmids were used. Both luciferase reporters (*pDefense-Eluc* and *pUBI-Redf*) were included in all treatments. In wheat cv. Lee (*Yr7*), only the *AvrYr7* candidate was transfected; in cv. AvocetS (no *Yr7*), *AvrYr7* was co-transfected with the *Yr7* plasmid to assess *Avr/R* interactions. *AvrSr50* served as a negative control, and *AvrSr50/Sr50* as a positive control. For dosage-response assays, *AvrYr7* plasmid amounts ranged from 1000ng to 50ng.

Wheat (*Triticum aestivum*) plants of cv. Lee, Thatcher, Vixen, Morocco, AvocetS and *Triticum spelta* were grown for 8-10□d in a growth cabinet with 150□μmol□m^−2^ □s^−1^ light intensity and at 21°C with 16□h day length. Protoplasts were isolated from those young leaves, followed by dispensing the plasmid mixes into sterile 96-well V-bottom plates. Each well received 50 μL of freshly isolated protoplasts, followed by 50 μL PEG-Ca solution. After gentle mixing, samples were incubated for 10 min at room temperature. Transfection was stopped with 200 μL W5 solution, and plates were incubated under low light at room temperature for 16 h prior to luminescence measurement.

After 16 h incubation, W5 supernatant was carefully removed and protoplasts were lysed in 50 μL 1x cell lysis buffer (Promega, #E3971) for 15 min. Lysate (50 μL) was transferred to opaque white 96-well plates (Corning, #CLS3600). Baseline luminescence was first recorded without substrate (1000 ms integration, 0 ms settle time) using a Tecan Infinite 200Pro plate reader. Steady-Glo substrate (50 μL; Promega, #E2520) was then added and luminescence measured again. Signals were acquired sequentially without filters (total luminescence), then through Lumi Green (∼505–560 nm) filter for defense-inducible reporter (*pDefense-Eluc*) and Red NB long-pass (cut-on ∼600 nm) filter for constitutive normalization reporter (*pUBI-pRedf*) under the same acquisition settings. Both reporters use luciferin substrate but emit distinct spectra, enabling ratiometric normalization.

### Multiple Sequence Alignment and Analysis of AvrYr7 alleles

*AvrYr7* alleles in genomes of different isolates were identified with BLAST^68^. Multiple sequence alignments of nucleotide and amino acid sequences were performed in Geneious Prime v2025.1.3 using the Clustal Omega (1.2.2) algorithm^69^.

### Naming of *AvrYr7* alleles and loci

We adopted an extended *Avr* allele nomenclature as introduced by Spanner et al., 2026 for *P. graminis* f. sp. *tritici*^6^. *Avr* variants were assigned a two-digit identifier based on their encoding unique protein sequence, e. g. AvrYr7-01, avrYr7-02. The suffix differentiates proteins variants with synonymous changes in the coding sequencing. We added a prefix with “*pN*” to differentiate regulatory promoter sequences with demonstrated phenotypic effects, such as infection or expression phenotypes, as exemplified by *p1-AvrYr7-01* and *p2-AvrYr7-07*. Where *p2* is associated with reduced expression relative to *p1*.

### RNAseq expression analysis of AvrYr7

Transcript abundance of AvrYr7 alleles was quantified using a time course of long-read ONT cDNA data for three isolates representative of distinct lineages. This included the previously published dataset for Pst104E which comprises ungerminated urediniospores (UG) and infected wheat leaf tissues at 4, 6, 8, 10 and 12 days post infection (dpi) (PRJNA1195871)^9^, as well as newly generated datasets for Pst239E (PstS10) and Pst134E (PstS1), each comprising UG, 8 and 10 dpi. All conditions across those isolates were sampled with four biological replicates. Long-read cDNA reads were trimmed using Porechop_ABI ^70^ and then mapped against each isolate’s phased dikaryotic assembly^16,47^ with minimap2^60^ in splice mode (-ax splice -ub -G 3000 --secondary=no) (. Only mappable reads were retained to filter out host transcripts. To obtain CDS annotations from Pst239E and Pst134E, we projected Pst104E gene annotations onto each haplotype assembly using Liftoff (-chrom -copies)^71^. *AvrYr7* projections were examined to confirm valid ORF and the absence of extra copies. Transcript abundance was quantified from splice alignments and CDS annotations using bambu^72^, and the resulting gene-level read counts were concatenated into a matrix for each isolate. Count matrices were analysed with DESeq2^73^, first to generate PCA and Cook’s distance plots for outlier detection, followed by median-of-ratios normalisation. Expression differences between each isolate’s allele pair were tested using Mann-Whitney *U* test.

### AlphaFold modelling

Multiple amino acid sequence alignment of the AvrYr7 variants was generated using ESPript^74^. Domain predictions of AvrYr7 variants were performed through the NCBI conserved domain search^75^. The structures of AvrYr7 variants, and AvrYr7-01 and Yr7 interaction complex, were modelled using AlphaFold3^35^. Structural alignments of AvrYr7 variants were performed using USalign. Protein homologous structure search was performed using DALI^76^ against Protein Data Bank. Protein structures are visualised using Pymol^77^.

## Supporting information

CDS sequences of alleles (fasta file)

AA sequences of alleles (fasta file)

1.Gene and promoter sequences of alleles (fasta file)

## Extended Data Figures

**S1: Defense reporter activation of *AvrYr7-01* in Thatcher and Vixen**. Defense signaling in protoplasts of *Yr7* containing wheat cv. **(a)** cv. Thatcher and **(b)** cv. Vixen was activated by *AvrYr7-01* and not by *avrYr7-02. AvrSr50* and *AvrSr50:Sr50* served as negative and positive controls respectively.

**S2: No defense activation of *AvrYr7-01* in *Triticum spelta* and *Triticum aestivum* cv. Morocco and AvocetS. (a)** *AvrYr7-01* and *avrYr7-02* did not induce the defense signaling in protoplasts of wheat having mismatched Yr gene, i.e., *Yr5* containing *T. spelta*. **(b)** Expression of *AvrYr7-01* and *avrYr7-02* in protoplasts of wheat cv. lacking any major Yr genes including *Yr7*, i.e., Morocco and AvocetS, also did not induce the defense signaling. *AvrSr50* and *AvrSr50:Sr50* served as negative and positive controls respectively.

**S3: Infection phenotypes in isolates from different *Pst* lineages. (a)** Representative infection phenotypes of *Puccinia striiformis* f. sp. *tritici* (*Pst*) isolates on cv. Morocco and cv. Lee (*Yr7*). Leaves of wheat cv. Morocco (susceptible control) and cv. Lee carrying the resistance gene *Yr7* were inoculated with the *Pst* isolates DK214_12 (*PstS0*), DK267_17 (*PstS10*), and RW10_15 (*PstS1*). Infection types (ITs) were scored visually on the second leaves 18 days after inoculation using a 0–9 scale where ITs 0–6 were classified as resistant and 7–9 as susceptible. All three isolates produced abundant uredinia on Morocco (IT 7-8). On cv. Lee, DK214_12 induced an incompatible chlorotic/necrotic response with little or no sporulation (IT 2-3), whereas DK267_17 and RW10_15 produced compatible phenotypes with abundant uredinia (IT 7). Scale bar = 5 mm.Representative infection phenotypes of *Puccinia striiformis* f. sp. *tritici* (Pst) isolates on cv. Morocco and cv. Lee (*Yr7*). Leaves of wheat cv. Morocco (susceptible control) and cv. Lee carrying the resistance gene *Yr7* were inoculated with the *Pst* isolates DK214_12 (*PstS0*), NL72067 (*PstS0*), RW10_15 (*PstS1*), DK09_11 (*PstS7*), DK02d_12 (*PstS8*), and DK69_15 (*PstS13*). Infection types (ITs) were scored visually on the second leaves 18 days after inoculation using a 0–9 scale, where ITs 0–6 were classified as resistant and 7–9 as susceptible^26,52,53^. All isolates produced abundant uredinia on Morocco (IT 7-8). On cv. Lee, DK214_12 induced an incompatible chlorotic/necrotic response with little or no sporulation (IT 2-3), whereas the remaining isolates produced compatible phenotypes with abundant uredinia (IT 7). Scale bar = 5 mm.

**S4: Variability of AvrYr7 alleles. (a)** schematic and **(b)** detailed protein alignments highlighting length differences, the variable C-terminal region, and key differences. Alignments are sorted by sequence similarity.

**S5: Non-suppression of *AvrYr7-07* recognition by *avrYr7-02* in wheat cv. Lee**. A similar defense induction was observed by *AvrYr7-01* and *AvrYr7-07* separately and when co-expressed with *avrYr7-02* in Lee protoplasts.

**S6: Absence of dosage dependent effect of *AvrYr7-07* as compared to *AvrYr7-01* in wheat cv. Lee**. *AvrYr7-01* and *AvrYr7-07* exhibited comparable defense activation in Lee protoplasts at varied plasmid amounts (1000ng – 50ng) with lower activation at 100 and 50ng.

**S7: Alignment of DNA sequences of all alleles including promoter region (1kb upstream of start codon)**. Black lines indicate differences between alignments.

## Extended Data Tables

**TableS1:** List of shared mutated candidate effector genes in DK06_00 and NL72067.

**TableS2:** Overview of isolates, alleles and virulence phenotypes.

**TableS3:** RNA-seq normalised expression values of alleles.

**TableS4:** Plasmids generated and used in this study.

## Supplemental files

1. Gene and promoter sequences of alleles (fasta file)
2. CDS sequences of alleles (fasta file)
3. AA sequences of alleles (fasta file)

## Acknowledgements

This work was supported by computational resources provided by the Australian Government through the National Computational Infrastructure (NCI) under the ANU Merit Allocation Scheme.

We acknowledge the contribution of the Plant Pathogen ‘Omics Initiative consortium in the generation of data used in this publication. The Initiative is supported by funding from Bioplatforms Australia, enabled by the Commonwealth Government National Collaborative Research Infrastructure Strategy (NCRIS). The authors acknowledge the provision of computing and data resources provided by the Australian BioCommons Leadership Share (ABLeS) program. This program is cofunded by Bioplatforms Australia, enabled by the Commonwealth Government National Collaborative Research Infrastructure Strategy (NCRIS), the National Computational Infrastructure and Pawsey Supercomputing Research Centre^78^.

## Data availability

Raw genome sequencing data of isolates DK06_00 (PstS0) and NL72067 (PstS0) and long-read cDNA reads for isolates Pst134E (PstS1) and Pst239E (PstS10) have been deposited on NCBI under Bioproject PRJNA1457771.

Genome assemblies of isolates DK06_00 (PstS0) and NL72067 (PstS0) and an updated version of the previously published Pst104E (PstS0) assembly and annotation have been deposited at zenodo (doi: 10.5281/zenodo.19804079).

Previously published genomes for Pst239E (PstS10) and PstDK09_11 (PstS7) can be found at https://zenodo.org/records/18396413. Genome assembly and RNA-seq of Pst104E have been previously published and are deposited at https://zenodo.org/records/14885411 and under NCBI Bioproject PRJNA1195871. The previously published Pst134E assembly can be accessed via NCBI Bioproject PRJNA749614.

