## Supplementary figures and images for "The variable wheat stripe rust effector *AvrYr7* evades *Yr7* recognition through sequence and expression polymorphisms"

### Extended Data Fig. 1

**Extended Data Fig. 1**

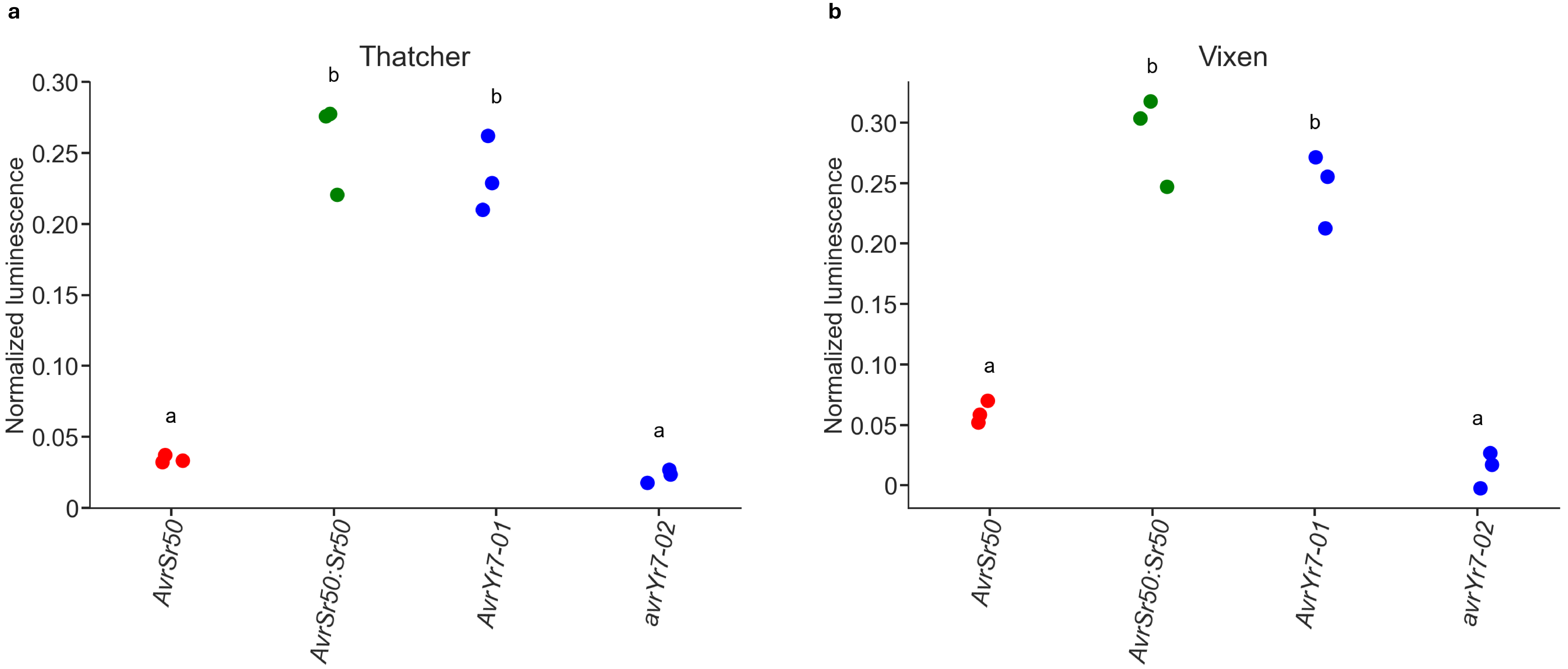

### Extended Data Fig. 2

Extended Data Fig. 2

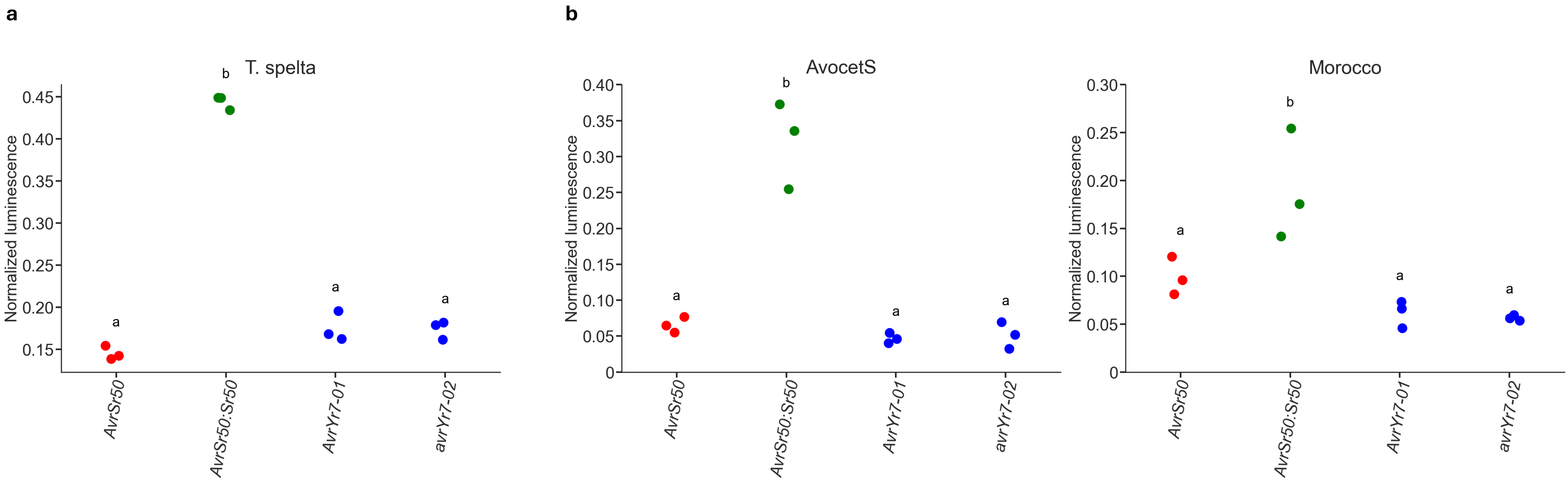

### Extended Data Fig. 3

**Extended Data Fig. 3**

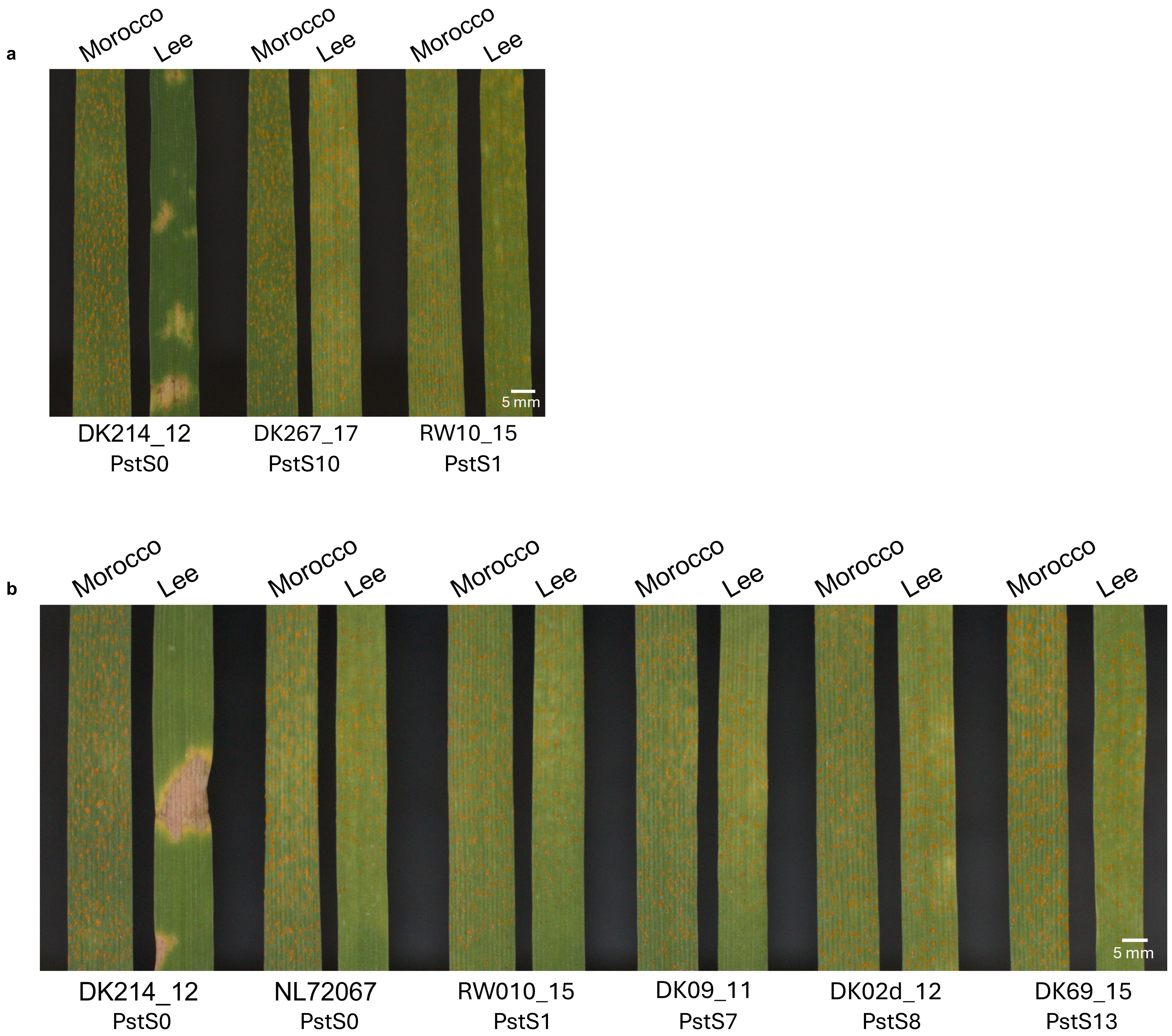

### Extended Data Fig. 4

**Extended Data Fig. 4**

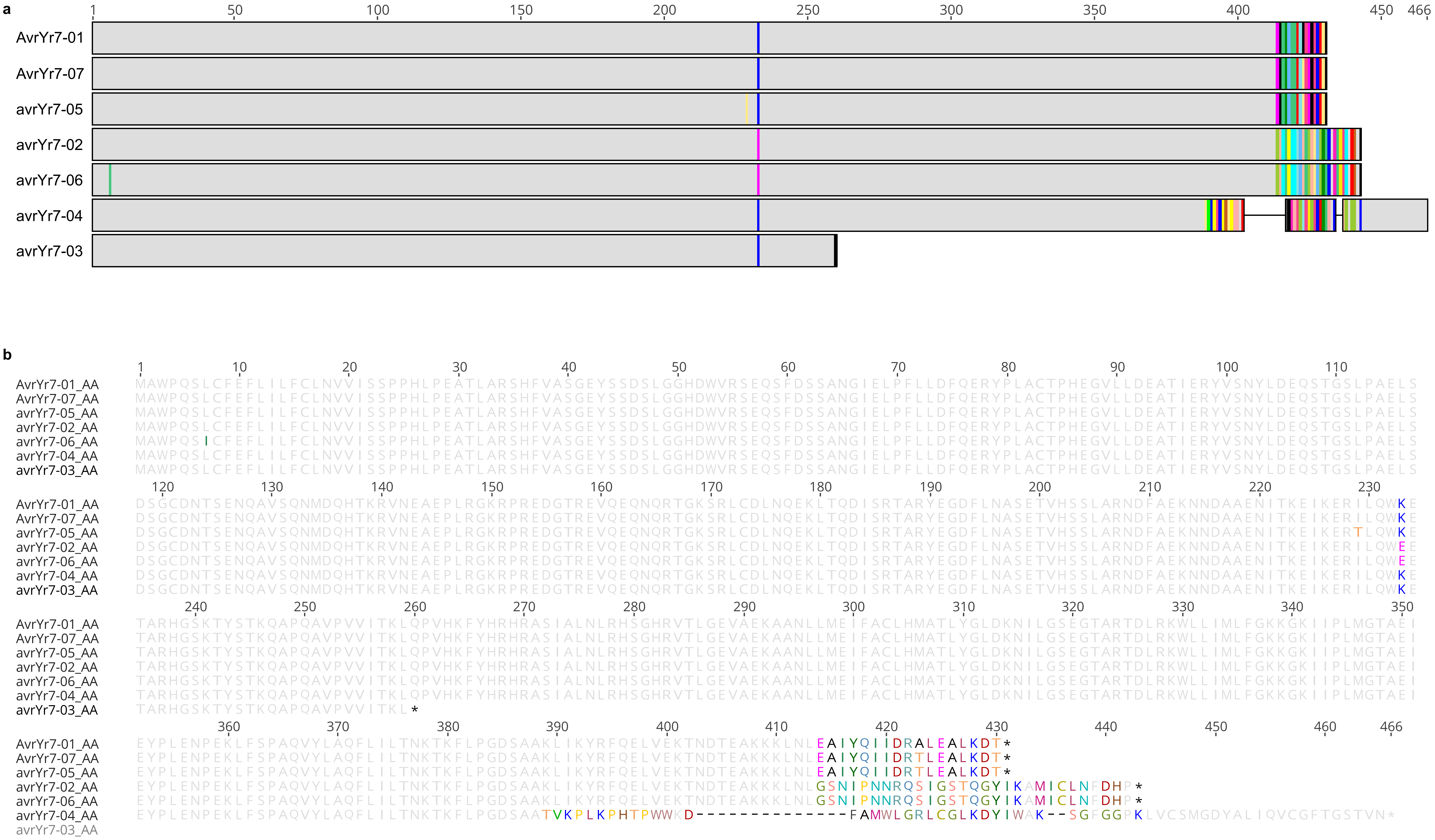

### Extended Data Fig. 5

**Extended Data Fig. 5**

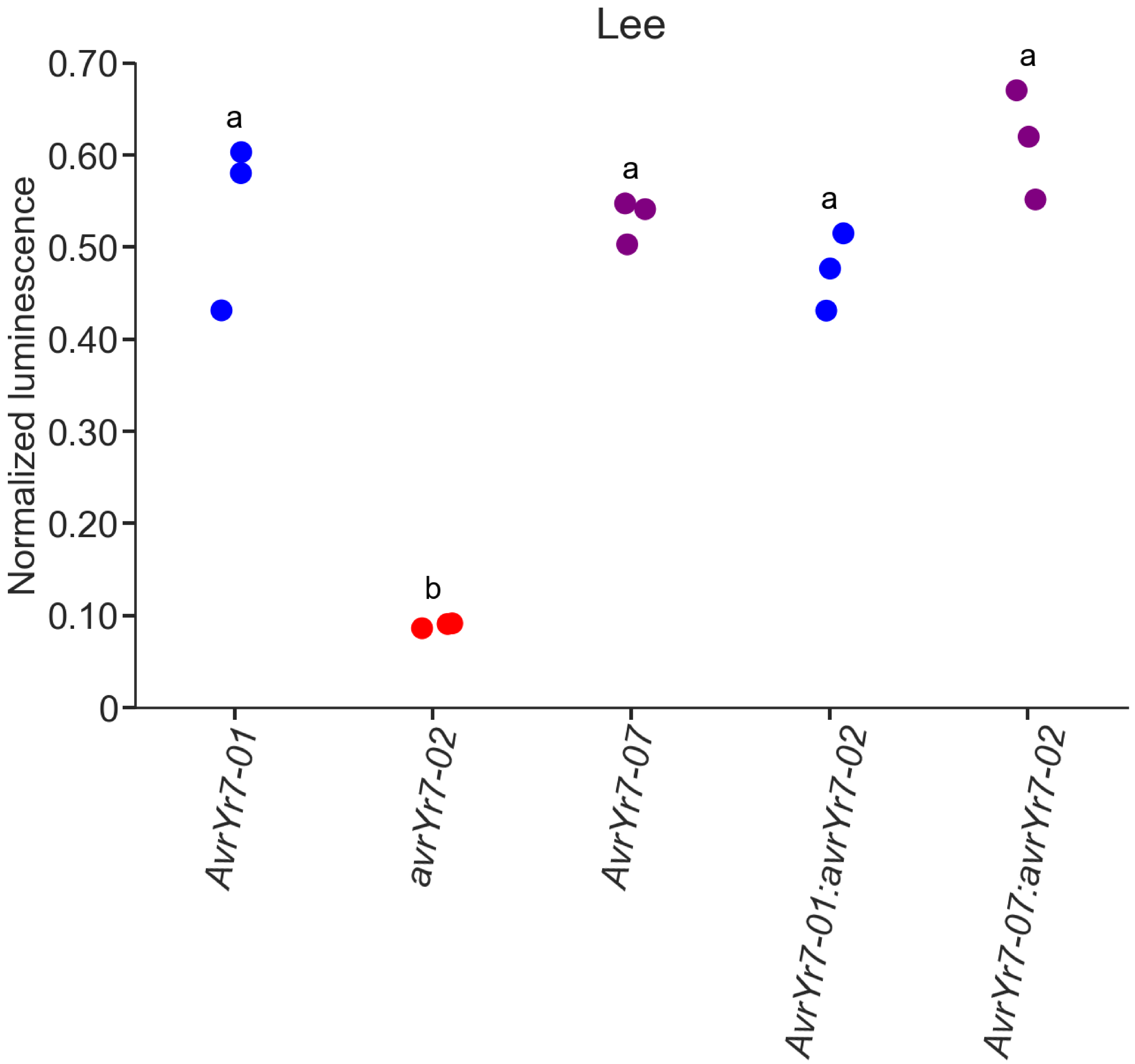

### Extended Data Fig. 6

**Extended Data Fig. 6**

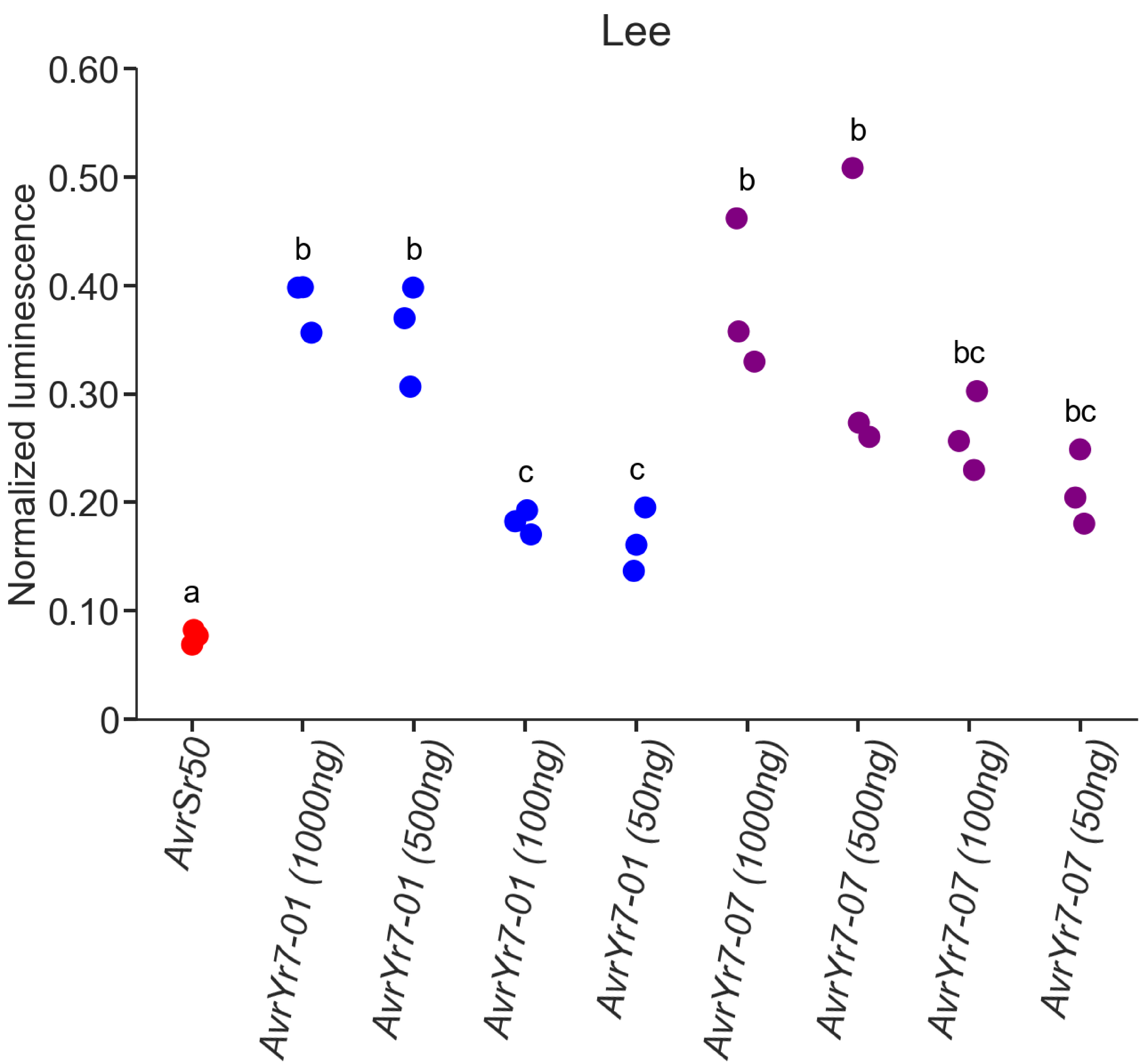

### Extended Data Fig. 7

**Extended Data Fig. 7**

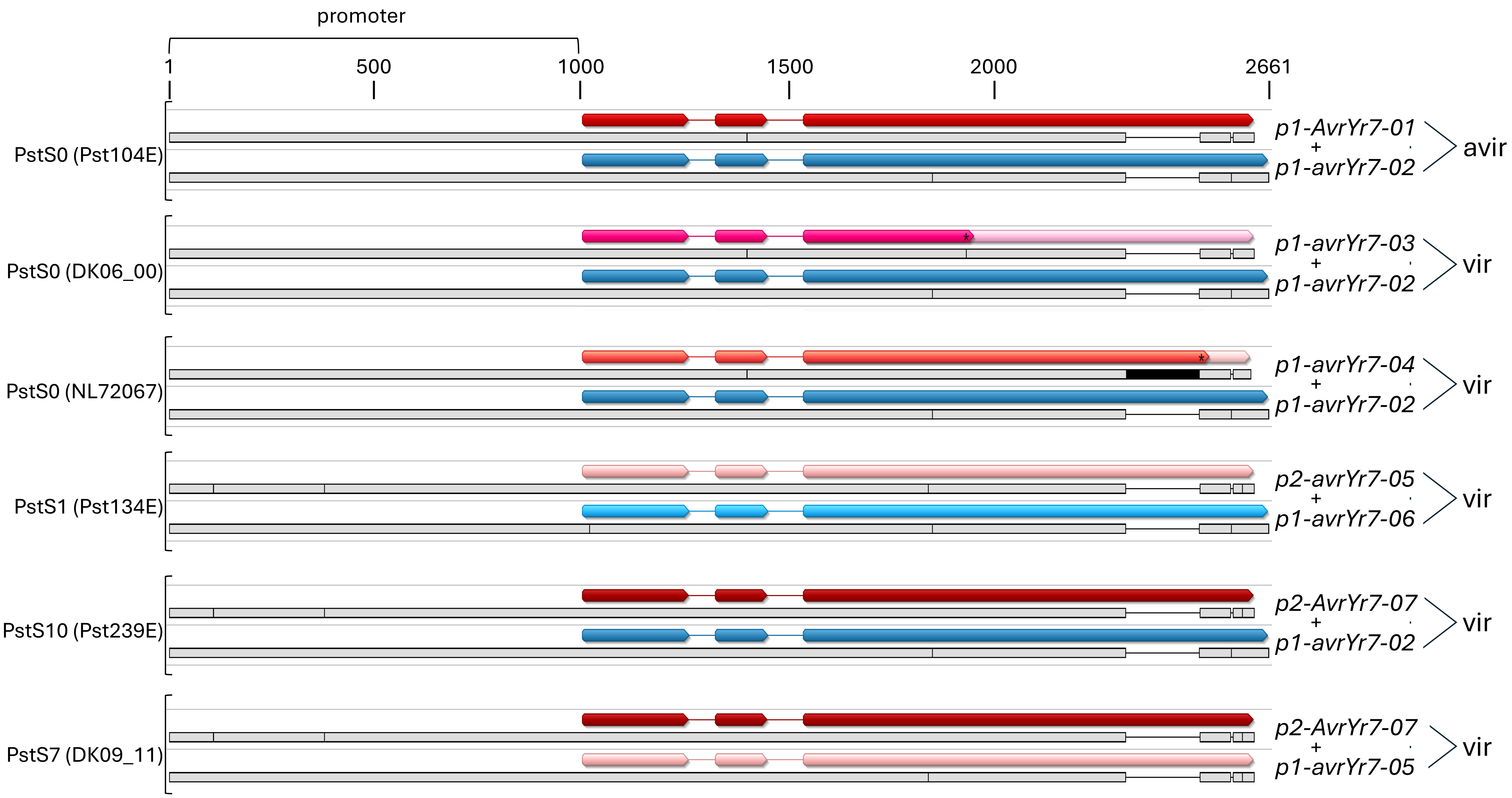
